# Heterologous sarbecovirus receptor binding domains as scaffolds for SARS-CoV-2 receptor binding motif presentation

**DOI:** 10.1101/2023.08.21.554179

**Authors:** Blake M. Hauser, Maya Sangesland, Evan C. Lam, Kerri J. St. Denis, Maegan L. Sheehan, Mya L. Vu, Agnes H. Cheng, Alejandro B. Balazs, Daniel Lingwood, Aaron G. Schmidt

## Abstract

Structure-guided rational immunogen design can generate optimized immunogens that elicit a desired humoral response. Design strategies often center upon targeting conserved sites on viral glycoproteins that will ultimately confer potent neutralization. For SARS-CoV-2 (SARS-2), the surface-exposed spike glycoprotein includes a broadly conserved portion, the receptor binding motif (RBM), that is required to engage the host cellular receptor, ACE2. Expanding humoral responses to this site may result in a more potently neutralizing antibody response against diverse sarbecoviruses. Here, we used a “resurfacing” approach and iterative design cycles to graft the SARS-2 RBM onto heterologous sarbecovirus scaffolds. The scaffolds were selected to vary the antigenic distance relative to SARS-2 to potentially focus responses to RBM. Multimerized versions of these immunogens elicited broad neutralization against sarbecoviruses in the context of preexisting SARS-2 immunity. These validated engineering approaches can help inform future immunogen design efforts for sarbecoviruses and are generally applicable to other viruses.

## Introduction

The emergence of SARS-CoV-2 (SARS-2) highlighted the need to leverage existing, as well as new, vaccine platforms to counter novel pathogens [1, 2]. Current SARS-2 vaccines have predominantly used a prefusion stabilized version of its surface glycoprotein, spike, which is effective against both the original USA-WA1/2020 strain as well as subsequent variants (*e*.*g*., B.1.1.7, BA.5) [3, 4]. This general approach for stabilizing a viral glycoprotein was first demonstrated for influenza hemagglutinin by introducing non-native cysteines or prolines to covalently “staple” or stabilize, respectively, the prefusion state [5, 6]. Prefusion stabilization was extended to viral glycoproteins from multiple viruses including other coronaviruses, respiratory syncytial virus, HIV, measles, Ebola, and Marburg [2, 7-11]. This illustrates how structural knowledge informs rational immunogen design for next-generation viral vaccines against emerging pathogens [12].

An additional approach used in structure-guided immunogen design is epitope scaffolding or “resurfacing” where a viral epitope is grafted on a comparatively unrelated protein scaffold [12-16]. These scaffolds are often based on antigenically distinct but structurally-related viral proteins, as well as *de novo* designed scaffolds; the latter approach represents a considerable design hurdle especially when the grafted epitope is complex or conformation-specific [15, 17, 18]. However, when using related viral proteins as potential scaffolds, sequence identity outside the grafted epitope may influence immune focusing effect by eliciting memory responses to additional epitopes within the scaffold. In the design process, this concern must be weighed against the engineering hurdles in a *de novo* designed scaffold that accurately recapitulates the conformation of the grafted epitope.

This engineering approach is particularly useful in context of prior humoral immunity as the designed immunogens, in theory, should recall potential memory responses imprinted to the displayed epitope. Indeed, prior humoral responses are often strain-specific and are recalled at the expense of more broadly protective responses [19-21]. However, *in vivo* evaluation of engineered immunogens often occurs in naïve animal models, which fails to recapitulate complex immune histories observed in humans [22-30]. While it is not possible to accurately encompass the entire diversity of immune histories in animal models, it remains critical to test immunogens within the context of some preexisting immunity. This is particularly necessary when the goal is to elicit broadly reactive responses that may be subdominant relative to other more strain-specific responses in the imprinted repertoire. Thus, it is necessary to understand the parameters that govern the antigenicity and immunogenicity of the engineered immunogens to maximize recall of broadly protective responses to the grafted epitopes.

Here, we used our resurfacing approach to graft the receptor binding motif (RBM) from SARS-2 onto antigenically distinct, but structurally related sarbecovirus receptor-binding domains (RBDs). The RBM is a known target of broadly neutralizing antibodies and eliciting humoral responses to this epitope may provide broad spectrum sarbecovirus immunity. We specifically selected sarbecovirus RBD scaffolds with increasing antigenic diversity, relative to SARS-2, to interrogate the relationship between varying antigenic distance and the subsequent humoral response towards the grafted, RBM epitope. Importantly, we performed these studies in the murine model previously immunized with the SARS-2 spike to simulate imprinted humoral immunity. Briefly, we identified additional sarbecovirus RBD scaffolds that accepted the SARS-2 RBM graft to maximize antigenic distance between the scaffolds but determined that further antigenic distance is required for RBM immune focusing. These data show the importance of antigenic diversity in scaffold selection and how further iterations of this design approach may lead to next-generation sarbecovirus vaccines. Moreover, it serves as a generalizable strategy for other pathogens where potential imprinted subdominant responses need to be recalled and expanded.

## Results

Both wildtype sarbecovirus and engineered RBDs elicit a broadly neutralizing antibody response [14, 24]. For the former, the response focused on portions of the RBD outside of the RBM; this was rather expected given that the non-RBM portions of the RBDs are conserved across the different sarbecoviruses [14]. Grafting the conserved SARS-2 RBM onto heterologous sarbecovirus scaffolds would potentially alter this pattern, as the grafted RBM would be a single consistent, conserved epitope, relative to the remainder of the RBD. Thus, using multiple heterologous, antigenically distinct, sarbecovirus scaffolds “resurfaced” with the SARS-2 RBM would immune focus to this grafted epitope.

We identified 20 distinct heterologous sarbecovirus RBDs to serve as potential scaffolds for the SARS-2 RBM (**Fig. 1A**). These scaffolds were selected from clade 2 of the sarbecovirus subgenus and are more antigenically distinct than our previously used scaffolds based on the SARS-CoV-1 (SARS-1) and WIV-1 sarbecoviruses [14]. The clade 2 members were predominantly isolated from bats in southeast Asia, and these viruses are reported not to use ACE2 as a host receptor [31, 32].

**Figure 1.**
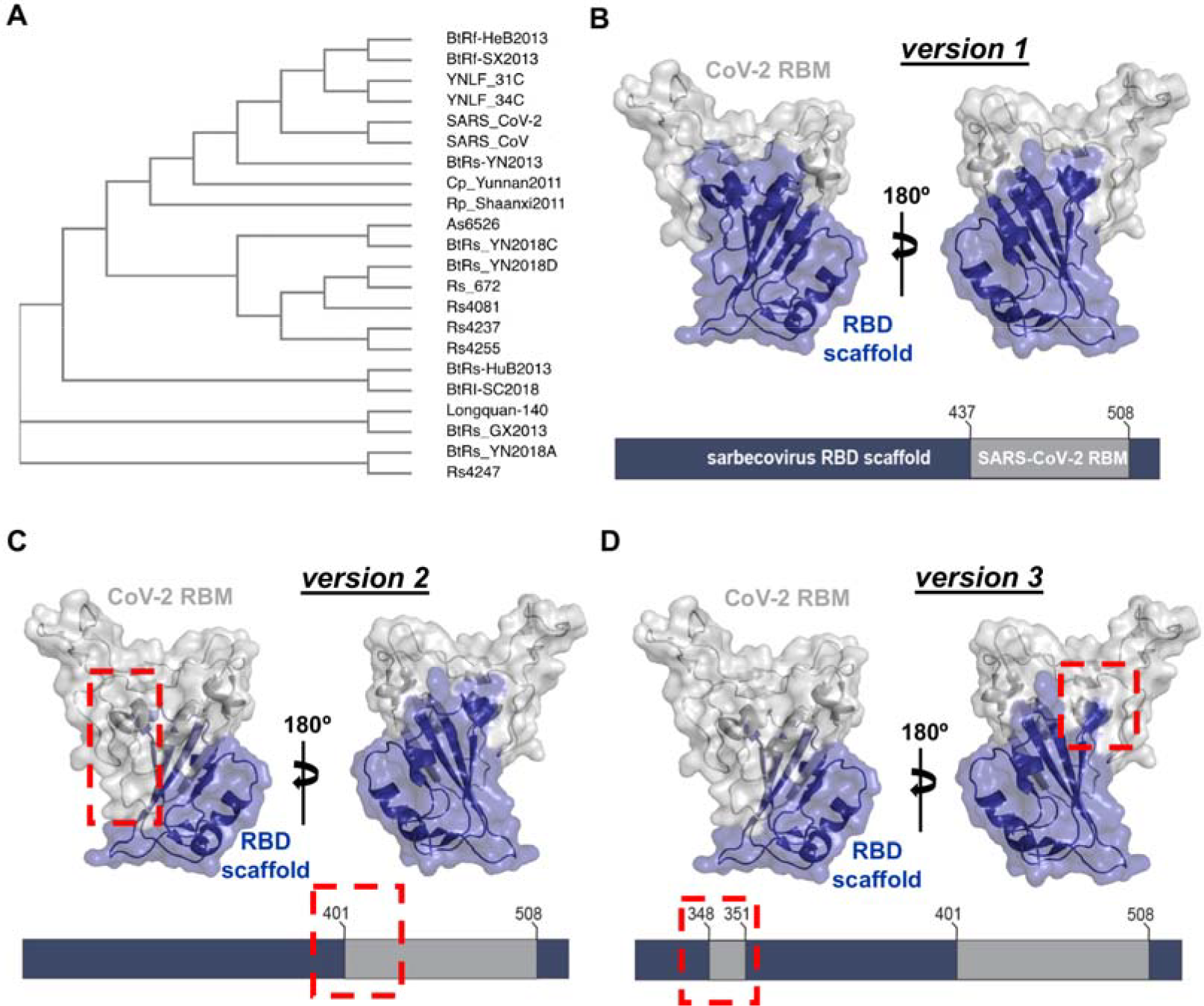
Selection of scaffolds and design. **(A)** Phylogenetic tree of twenty selected receptor binding domains used in this study; SARS-CoV and SARS-CoV-2 are included for reference. **(B)** Design schematic of versions 1, 2, and 3 resurfaced constructs, depicting RBM grafts (grey) on heterologous coronavirus scaffolds (indigo). Red dashed boxes indicate differences relative to version 1.

To graft the SARS-2 RBM onto these scaffolds, we used three different versions (v). The first, “v1”, used identical RBM boundaries as our initial resurfaced “rs”, rsSARS-1 and rsWIV1 constructs: residues 437-507 (SARS-2 spike numbering) (**Fig. 1B**) [14]. A second, v2, used a more extensive RBM graft spanning residues 401-507 which was shown previously to confer ACE2 binding (**Fig. 1C**) [31]. Based on additional structural analysis, our final v3 construct incorporated SARS-2 RBD residues 348-352 in context of v2 to potentially stabilize the SARS-2 RBM in context of the heterologous scaffold (**Fig. 1D**).

We first attempted to express the v1 constructs for 20 different clade 2 sarbecovirus RBDs. Of these 20 constructs, 5 could be recombinantly expressed in mammalian cells to varying degrees: rsBtRs-HuB2013, rsYNLF-34C, rsRs4081, rsRs4255, and rsBtRI_SC2018. However, further analysis by size exclusion chromatography (SEC) showed that these proteins were prone to aggregation. We then expressed the v2 and v3 constructs for these five constructs to determine if extending the graft and additional stabilizing residues would enhance expression and reduce aggregation. While all five v2 constructs had increased expression relative to v1, rsYNLF-34C had the highest level of expression and homogenous, monomeric protein was isolated via SEC (**Fig. 2**); all five v3 constructs expressed at a level between the v1 and v2 constructs.

**Figure 2.**
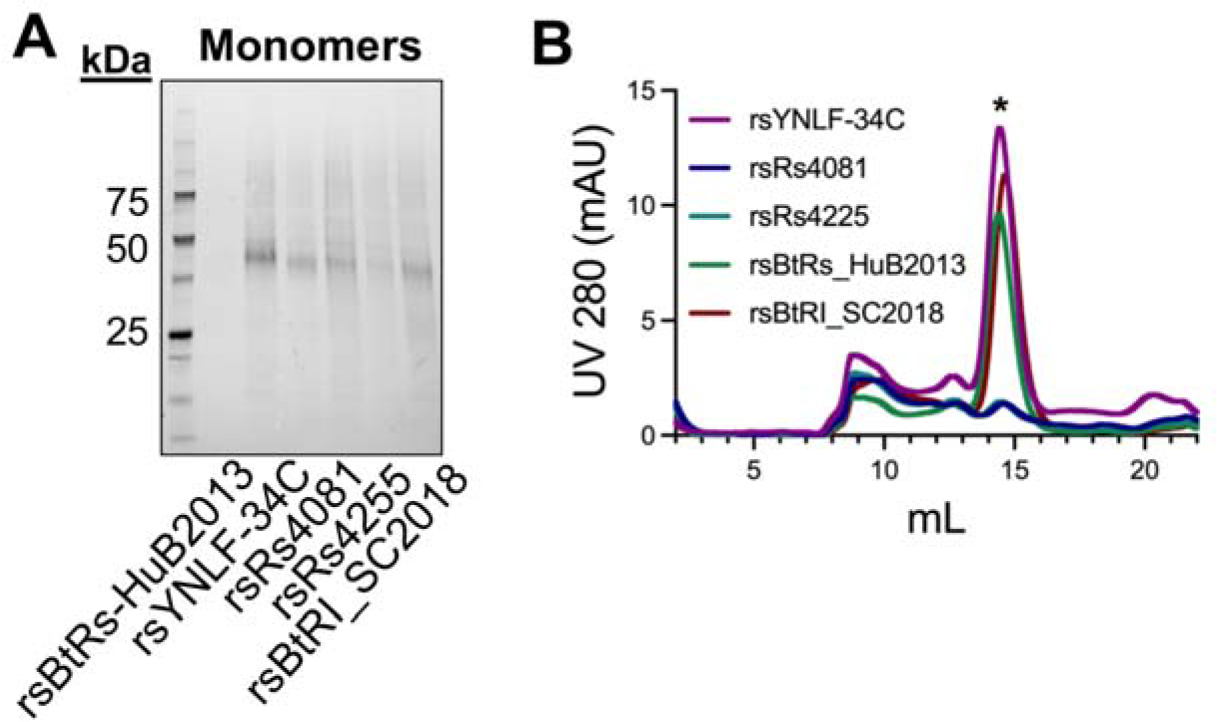
Resurfaced monomeric immunogen expression. **(A)** SDS-PAGE analysis of monomers under non-reducing conditions. **(B)** Representative size exclusion trace with (*) marking the monomeric constructs. Fractions in this peak were pooled for further characterization.

We next assessed rsYNLF-34C for binding to conformation-specific antibodies using biolayer interferometry (BLI). It bound SARS-2 RBM-directed antibody, B38, with comparable affinity to the wild-type SARS-2, with a monomeric-monomeric K_*D*_ of ∼1.4 μM (**Fig. 3A**) [33]; however, rsYNLF-34C did not have any detectable binding to CR3022 or S309 Fabs which bind outside the RBM [34, 35] (**Fig. 3A**). This lack of sequence conservation in the non-RBM regions of rsYNLF-34C relative to the SARS-2 RBD suggests that such a construct may enhance RBM-focusing by minimizing cross-reactive epitopes elsewhere in the RBD; we thus chose to advance this construct to test this hypothesis.

**Figure 3.**
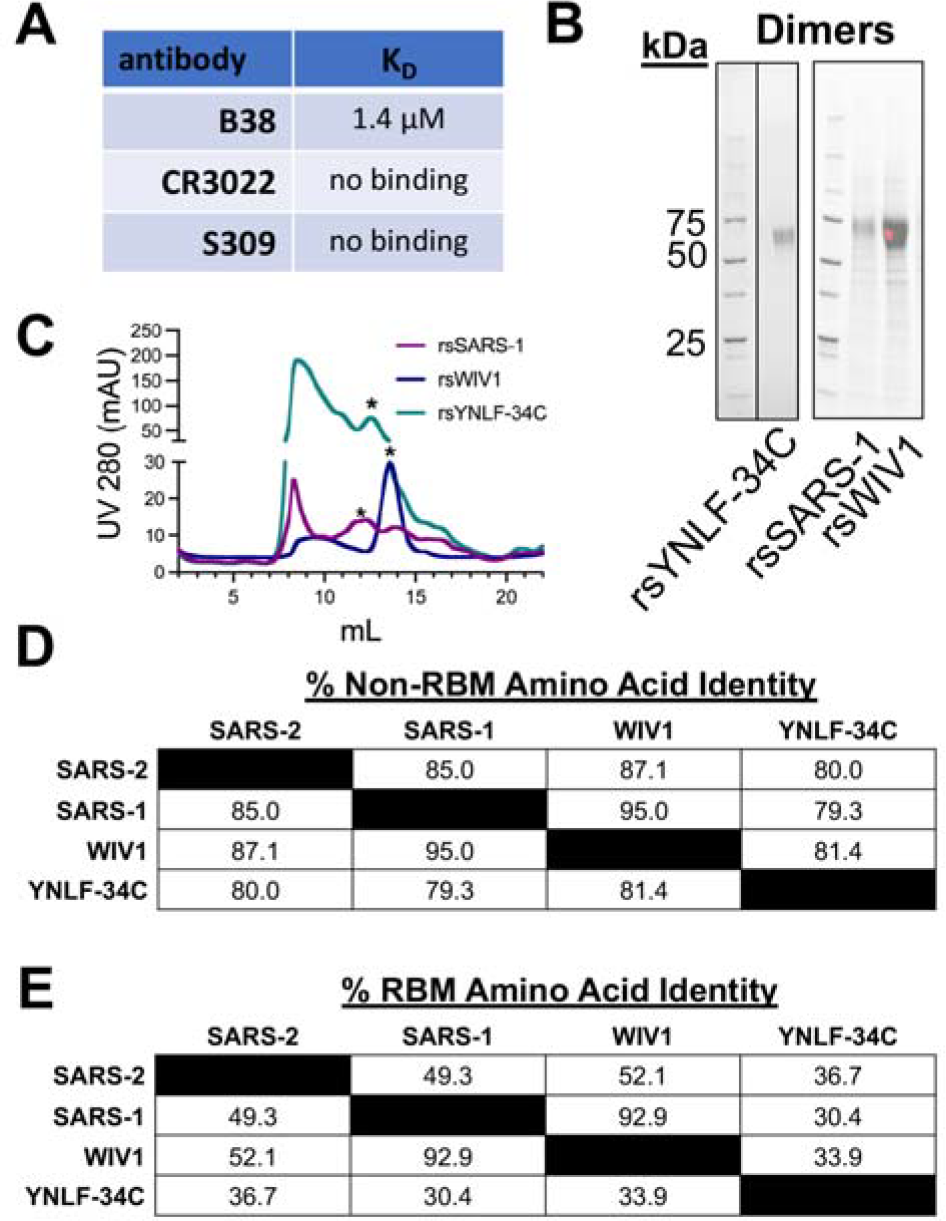
Resurfaced dimeric immunogen design and expression. **(A)** RBM-directed Fab B38 was used to confirm conformational integrity of the SARS-2 RBM grafted onto rsYNLF-34C. FAB2G sensors were used with immobilized Fab to measure binding via BLI to rsYNLF-34C at 10 μM, 5 μM, 1 μM, and 0.5 μM. Binding of CR3022 and S309 Fabs to rsYNLF-34C at 10 μM was also assayed, but no binding was detected. **(B)** SDS-PAGE analysis of dimers under non-reducing conditions. **(C)** Representative size exclusion trace with (*) marking the dimeric constructs. Fractions in this peak were pooled for use as immunogens. **(D)** Pairwise comparisons of amino acid sequence identity in the non-RBM portions of selected sarbecovirus RBDs. **(E)** Pairwise comparisons of amino acid sequence identity in the RBM portions of selected sarbecovirus RBDs.

We next engineered dimeric forms of rsYNLF-34C and our original rsSARS-1 and rsWIV1 immunogens to improve immunogenicity as previously described for wild-type coronavirus RBDs [22]. The rsWIV1 construct used the “end-to-end” dimerization approach previously described [22], while the rsSARS-1 and rsYNLF-34C dimers had a 13 amino acid glycine-serine linker between the two RBDs to enhance construct expression. These constructs were confirmed using SDS-PAGE analysis, and the dimeric species was isolated using SEC (**Fig. 3B, C**). The rsYNLF-34C, rsSARS-1, and rsWIV-1 scaffolds range in antigenic distance from ∼79.3% to ∼95.0% amino acid identity relative to SARS-2 within non-RBM portion of the RBD, while the amino acid identity ranged for each wild-type RBM ranged from ∼30.4% to ∼92.9% (**Fig. 3D, E**).

To test if increased antigenic distance between coronavirus RBD scaffolds would enhance SARS-2 RBM focusing, we designed an immunization regimen with two different cohorts (**Fig. 4**). The first included a cocktail of the rsSARS-1 and rsWIV1 dimers, which share 95.0% amino acid sequence identity between the scaffolds in the non-RBM portion of the RBDs and referred to as antigenic distance “low” (AgDist_*low*_). The second, included rsYNLF-34C and rsWIV1 dimers, which have 81.4% amino acid and referred to as AgDist_*high*_.

**Figure 4.**
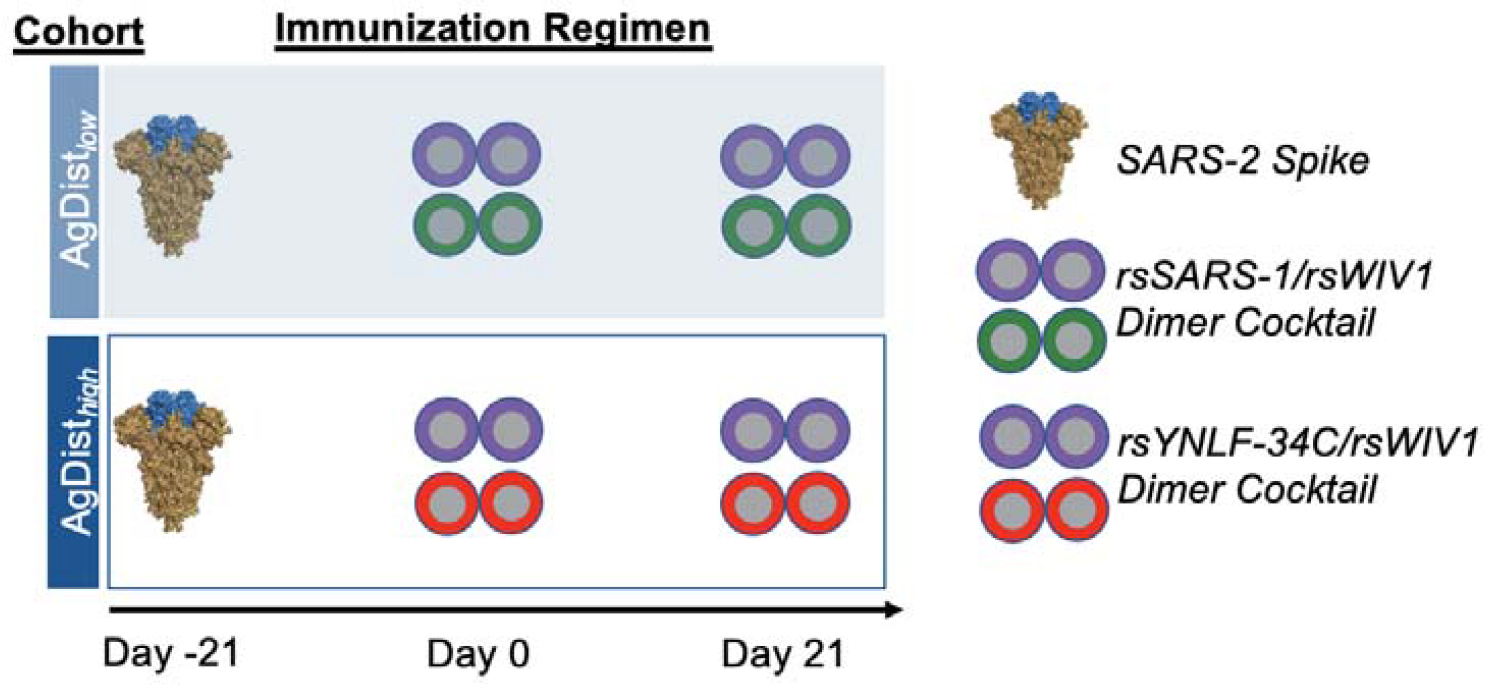
Antigenic distance investigation immunizations. Schematic of immunization regimens. Two immunization cohorts (n=5 mice each) were primed with SARS-2 two-proline-stabilized (2P) spike protein on day 0 and then boosted with a cocktail of either the rsSARS-1 and rsWIV1 dimers (“AgDist_*low*_” cohort) or the rsWIV1 and rsYNLF-34C dimers (“AgDist*high*” cohort) at days 21 and 42.

We then immunized two cohorts of C57BL/6 mice that were first primed with recombinant SARS-2 spike to simulate preexisting immunity and then immunized and boosted with either AgDist_*low*_ or AgDist_*high*_ immunogens (**Fig. 5A**). Serum was collected at day 35, and ELISAs were performed against a variety of coronavirus proteins. To assess potential RBM focusing, we used a previously characterized SARS-2 construct (RBM^*hg*^), that has glycans placed across the RBM to abrogate binding of RBM-directed antibodies (*e*.*g*., B38) [14]. Neither cohort, however, showed a significant difference in titers against the wild-type SARS-2 RBD and the SARS-2 RBM^*hg*^ construct, indicating a lack of serum antibody response focusing to the SARS-2 RBM. This suggests that additional antigenic diversity between scaffolds may be required.

**Figure 5.**
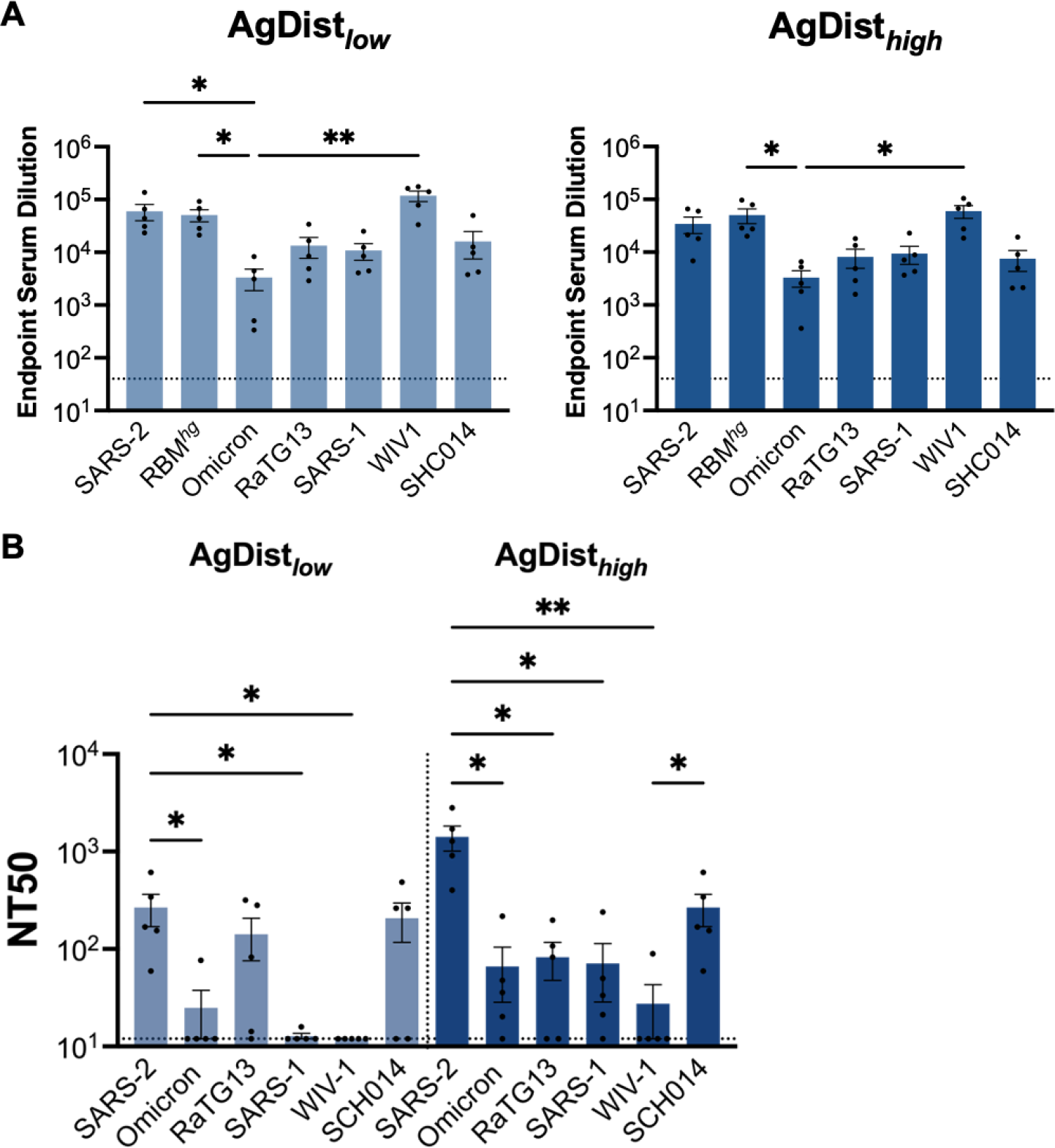
Antigenic distance investigation immunization serum reactivity. **(A)** Serum collected at day 35 was assayed in ELISA against different coronavirus antigens. **(B)** Pseudovirus neutralization was assayed against a panel of sarbecoviruses, ordered here based on spike amino acid sequence similarity to SARS-2. Statistical significance for **(A, B)** was determined using Kruskal-Wallis test with post-hoc analysis using Dunn’s test corrected for multiple comparisons (* = p < 0.05, ** = p < 0.01); non-significant comparisons not marked.

Despite the lack of observed RBM focusing, sera from the AgDist_*high*_ cohort showed detectable neutralization against all sarbecoviruses tested (**Fig. 5B**). This contrasts with the sera from the AgDist_*low*_ cohort, which showed limited neutralization against the SARS-2 Omicron variant, SARS-1, and WIV1 pseudoviruses (**Fig. 5B**). Immune focusing to conserved RBM epitopes has been previously shown to confer potent neutralization across a broad range of sarbecoviruses, indicating that there may be some level of immune focusing in the AgDist_*high*_ cohort even though this difference is not readily detected by ELISA [14].

## Discussion

This study builds upon our previous work that combined resurfacing and hyperglycosylation protein engineering approaches for potential immune focusing to conserved epitopes [14]. In that study, the SARS-1 and WIV-1 scaffolds had rather high sequence identity relative to each other as well as to SARS-2. Here, we used our resurfacing approach to identify additional sarbecovirus RBD scaffolds that could present the SARS-2 RBM to further maximize antigenic distance between the scaffolds. It appears, however, that further increasing this distance may still be required to achieve immune focusing. Additional investigations will be required to interrogate the exact scaffold design parameters required to achieve successful immune focusing to the grafted RBM. The extent to which the parameters determined for this resurfacing approach and scaffold identification apples to grafted epitopes from other viral epitopes, such as the influenza receptor binding site, is also of interest.

We note, however, that despite a detectable SARS-2 RBM focusing within our assay, the AgDist_*high*_ cohort did show improved neutralization breadth in the context of prior SARS-2 spike imprinting. This suggests that these the resurfaced immunogens and approach more broadly may serve as a foundation for next-generation pan-sarbecovirus vaccines by preferentially increasing the representation of the SARS-2 RBM within candidate immunogens. Broadly neutralizing, RBM-directed antibodies have also been characterized by others [32, 36-38]. Collectively, these findings show that simply immunizing with diverse sarbecovirus RBDs does not universally confer broad neutralization in the setting of SARS-2 spike imprinting. This is particularly interesting in the case of the WT Heterotrimer cohort, which utilizes the same three RBDs that we had previously shown to elicit a broadly neutralizing antibody response when a cocktail of homotrimeric RBDs was used as the boosting immunogen [14]. One possible explanation is that the orientation and subsequent positioning of the RBD components in a particular immunogen might affect the extent to which cross-reactive antibodies are elicited. Indeed, this has been suggested for other immunogens, in particular those displayed on various nanoparticles or DNA-based scaffolds [39, 40].

Immunization with a range of wild-type sarbecovirus RBDs also conferred broad neutralization both in naïve animals and in the context of SARS-2 spike imprinting [14, 24]. These broadly neutralizing antibodies preferentially target conserved epitopes on the SARS-2 RBD, with most contacts falling outside the RBM [14, 41]. Similar findings have been demonstrated for influenza after immunization with diverse hemagglutinins [39]. We recently showed that a similar resurfacing approach for influenza showed that grafting the complex, conformation-specific, hemagglutinin receptor binding site onto multiple antigenically distinct scaffolds, could preferentially expand responses to this site [42]. Thus, our study here provides further support, with further design iterations required, that this principle of “epitope enrichment” may be broadly applicable across virus families and could be leveraged to design immunogens that focus to any enriched epitope of interest.

The structure-guided, rational immunogen design approaches implemented in this study have the potential to inform future pan-sarbecovirus vaccine efforts. Additionally, these findings provide some mechanistic insight into parameters required to influence the humoral immune response towards desired epitopes.

## Methods

### Protein Expression and Purification

Dr. Jason McLellan at the University of Texas, Austin generously shared the SARS-2 spike plasmid, which contained a Foldon trimerization domain and C-terminal HRV 3C-cleavable 6xHis and 2xStrep II tags. RBD designs were based on the following sequences: SARS-2 RBD (Genbank MN975262.1), SARS-1 RBD (Genbank ABD72970.1), WIV1 RBD (Genbank AGZ48828.1), RaTG13 RBD (Genbank QHR63300.2), SHC014 RBD (Genbank QJE50589.1). Codon optimization was performed using Integrated DNA Technologies, and gblock constructs were purchased and cloned into the pVRC expression vector. Proteins included a C-terminal HRV 3C-cleavable 8xHis tag, as well as SBP tags in the monomeric and dimeric RBDs and a previously published hyperglycosylated, cystine-stabilized trimerization tag in the trimeric RBDs [14].

Constructs were expressed using Expifectamine transfection reagents in Expi293F cells (ThermoFisher) according to the manufacturer’s protocol. After 5-7 days, transfections were harvested, and supernatants were purified via immobilized metal affinity chromatography using Cobalt-TALON resin (Takara). Eluted proteins were further purified by size-exclusion chromatography using a Superdex 200 Increase 10/300 GL column (Cytiva) in PBS (Corning). Relevant fractions were pooled. Heterotrimeric proteins were further purified using anti-FLAG resin (Genscript) and eluted with 160 μg/mL FLAG peptide (APExBIO) in Tris buffered saline followed by streptavidin resin (Pierce) eluted with 4 mM biotin (Sigma) in HEPES buffer.

HRV 3C protease (ThermoScientific) was used to remove purification tags prior to immunizations. Cleaved protein was re-purified using Cobalt-TALON resin and size exclusion chromatography to separate uncleaved protein, protease, and cleaved tags from cleaved protein.

### Immunizations

Immunizations were performed in C57BL/6 female mice aged 6-10 weeks (Charles River Laboratories). Each mouse received 20 μg of protein adjuvanted with 50% w/v Sigma Adjuvant System in 100 μL of inoculum via the intraperitoneal route [43]. Immunizations occurred at days-21, 0, and 21, with serum characterization occurring with day 35 samples. All experiments were conducted with institutional IACUC approval (MGH protocol 2014N000252).

### Serum ELISAs

Serum ELISAs were performed as previously described [14]. Briefly, plates were coated with 100 μL of protein at 5 μg/mL overnight at 4 °C, then blocked with 150 μL of 1% BSA in PBS-Tween for 1 hour at room temperature. Sera was added at a starting dilution of 1:40 with a serial 5-fold dilution and incubated for 90 minutes at room temperature. Plates were washed, and 150 μL of HRP-conjugated anti-mouse IgG (Abcam) was added at 1:20,000. After a 1-hour secondary incubation at room temperature, plates were washed and 150 μL of 1xABTS development solution (ThermoFisher) was added. Plates were developed for 30 minutes at room temperature before stopping with 100 μL of 1% SDS to read on a SpectraMaxiD3 plate reader (Molecular Devices) for absorbance at 405 nm.

### Pseudovirus Neutralization Assay

Serum pseudovirus neutralization assays were performed as previously described [14, 44]. Briefly, pseudotyped lentiviral particles were generated via transient transfection of 293T cells and tittered using 293T-ACE2 cells [45] and the HIV-1 p24CA antigen capture assay (Leidos Biomedical Research, Inc.). Assays were performed using a Tecan Fluent Automated Workstation and 384-well plates (Grenier), and sera were initially diluted 1:3 with subsequent serial 3-fold dilutions. Each well received 20 μL each of sera and pseudovirus (125 infectious units), and plates were incubated 1 hour at room temperature. 10,000 293T-ACE2 cells [45] in 20 μL of media with 15 μg/mL polybrene was added, and additional incubation occurred at 37 ° C for 60-72 hours. Cells were lysed [46], and luciferase expression was quantified on a Spectramax L luminometer (Molecular Devices). Neutralization percentage was calculated for each serum concentration by deducting background luminescence from cells-only wells and dividing by the luminescence of wells containing cells and virus. GraphPad Prism (version 9) was used to fit nonlinear regressions, and IC50 values were calculated for samples with neutralization values that were at least 80% at maximum serum concentration. Reciprocal IC50 values were used to obtain NT50 values.

## Acknowledgements

We thank Dr. Jason McLellan from University of Texas, Austin for the spike plasmid. We thank Nir Hacohen and Michael Farzan for the kind gift of the ACE2 expressing 293T cells. We acknowledge funding from NIH R01s AI146779 (AGS); AI174875 (ABB); AI155447, AI137057, and AI153098 (DL), a Massachusetts Consortium on Pathogenesis Readiness (MassCPR) grant (AGS), and P01 AI168347 (AGS); training grants: NIGMS T32 GM007753 (BMH); T32 AI007245 (JF); F31 Al138368 (MS); F30 AI160908 (BMH). ABB is supported by the National Institutes for Drug Abuse (NIDA) Avenir New Innovator Award DP2DA040254, the MGH Transformative Scholars Program as well as funding from the Charles H. Hood Foundation (ABB). This independent research was supported by the Gilead Sciences Research Scholars Program in HIV (ABB).

## Author contributions

Conceptualization, BMH, AGS; Methodology, BMH, MS, ECL, AGS; Investigation, BMH, MS, ECL, KJS, MLS, MLV, AHC; Writing – Original Draft, BMH and AGS; Writing – Review and Editing, all authors; Funding Acquisition, ABB, DL, AGS; Supervision, ABB, DL, AGS.;

## Declaration of interests

The authors have no competing interests to declare.

## Data availability

The datasets generated during and analyzed during the current study are available from the corresponding author on reasonable request.

## Notes

### Competing Interest Statement

The authors have declared no competing interest.

